# Theta-Phase Targeted Encoding Gates Memory Sensitivity

**DOI:** 10.64898/2026.10.29.762605

**Authors:** Antonia Raissle, Vincent van de Ven, Inge Leunissen, Zhou Fang, Giancarlo Valente, Parisa Naseri, Sanne Ten Oever

## Abstract

Theta phase has been proposed to play an important role in memory. It may gate encoding and retrieval, such that certain phases are more favorable for memory processing than others, or become part of the memory representation, such that reinstating the encoding phase benefits retrieval. To test and distinguish these competing accounts, we used a closed-loop EEG system, allowing us to control the phase at which items were presented at encoding and retrieval. In an associative memory task, visual targets were presented at predefined phases of the ongoing mid-frontal theta rhythm, thereby pairing individual memory items with specific encoding phases. During retrieval, targets appeared either at the same phase as during encoding or at an alternative phase. We show that theta phase during encoding predicted subsequent memory sensitivity, while finding no evidence that retrieval phase nor the match between encoding and retrieval phases influenced performance. Therefore, our findings are consistent with a gating account, in which the rhythmic state at the moment of encoding influences the efficacy of memory formation.

## Introduction

Brain oscillations reflect the cyclical variations in neuronal excitability of populations of neurons ^1,2^. These fluctuations influence the likelihood of neuronal firing, with the highest likelihood of firing occurring during phases of neuronal depolarization ^3,4^. Importantly, by modulating moment-to-moment excitability, oscillatory phase shapes the processing of incoming sensory information and can consequently influence behavior. For example, oscillatory phase at the onset of a stimulus has been shown to predict perceptual outcomes. This was shown in the visual domain, where evidence suggests that alpha phase predicts visual stimulus detection ^5,6^, as well as in the auditory domain, where theta and alpha phase have been related to sound detection and speech categorization ^7–9^.

A growing body of research investigates if phase-dependent processes observed in perception extend to human memory. This research has focused on theta-band oscillations (4 – 8Hz), which have been consistently implicated in episodic memory encoding and retrieval ^10–12^. It has been proposed that theta phase might act as a mnemonic gate, such that information processed during a particular phase of the theta cycle is encoded or retrieved more successfully and subsequently remembered better ^13–17^. On this account, theta phase can be seen as a *process*, that for example gates information processing. Other accounts emphasize theta’s role in representing information, proposing that theta forms part of a contextual code ^18^. In this view, the specific phase at which an item is encoded could become bound to its memory representation, and by that, itself forming part of what is stored ^9^. We refer to this as the *phase-as-representation* account, in which phase functions as an internal reference to which an item is tied. This is in line with the theta-gamma code hypothesis, in which the theta oscillation provides a reference frame within which individual memory representations are organized ^19,20^. Accordingly, memory representations can occupy different phases of the theta cycle defined by the phase at encoding ^9^.

The two accounts-phase as *process* or phase as *representation*-form testable predictions. If phase gates the encoding or retrieval process, memory strength should be influenced dependent on which phase it was encoded or retrieved respectively. On the other account, if phase forms part of the memory representation, memory should be facilitated whenever encoding and retrieval phase of an item match: re-encountering the encoding phase paired with the memory item at retrieval should reinstate part of the representation, which ultimately should cue recall. To date, however, no study has directly compared these competing accounts by directly testing their predictions. Distinguishing them requires memory items to be tested at both their matching and non-matching encoding-retrieval phases. Conventional electroencephalogram (EEG) does not allow the relationship between item and phase to be experimentally controlled, as phase-dependent effects can only be investigated by sorting trials post-hoc ^5,6,9,21^. Because endogenous theta phase fluctuates unpredictably, the same item is presented at different phases across repetitions. To overcome this constraints, previous studies have resorted to alternative methods to infer or manipulate phase, including sensory entrainment ^22–25^, transcranial alternating current stimulation ^26–28^, phase alignment between multisensory rhythmic stimuli during encoding ^29^, or phase-resetting approaches, in which stimulus-evoked resetting of ongoing oscillation allows to infer about rhythmic fluctuations in memory encoding ^13^. Although these approaches have provided important insight into frequency– and phase dependent processing, they rely on externally induced rhythms, that might not accurately reflect endogenous oscillatory theta dynamics involved in episodic memory processes.

To gain experimental control over encoding and retrieval phase of memory items, the current study used a real-time closed-loop EEG setup together with an associative memory task, in which participants formed associations between initially unrelated abstract visual cue-target pairs ^30^. Because mid-frontal theta has been repeatedly implicated in successful memory encoding ^16,31^, we leveraged this oscillatory signal as a basis for the real-time phase estimation. Using the closed-loop EEG system, endogenous mid-frontal theta oscillations were monitored in real-time to trigger target presentation at one of two predefined phases (–0.5π and 0.5π) of the theta cycle during encoding, thereby establishing a stable item-phase relationship. During the subsequent memory test, the previously established item-phase relationship was no longer maintained, such that each target could appear either on the phase which it was encoded on or at the alternative phase, with which it had not previously been associated. In other words, target presentation either reinstated the phase context established during encoding or violated it. This design can separate a *process* signature of theta phase ^13,14^ (main effect of encoding/retrieval phase) from a *representational* one ^9,19,20^ (encoding – retrieval phase interaction).

## Results

In the current study, 43 participants performed an associative memory task while undergoing closed-loop EEG (see Methods for inclusion criteria). The experiment consisted of three task periods: a passive exposure, a learning, and a testing period. During the passive exposure period, participants were presented with eight abstract visual cue-target pairs, in which the cue and the target were presented sequentially. Each pair was shown twice. These pairs, randomly assigned to each participant, served as the participant’s matching pairs. Importantly, each pair was assigned to one of two pre-defined mid-frontal theta phases (–0.5π or 0.5π; further referred to as Learned phase). We randomized the assignment of phases and stimulus pair across participants, with four pairs at each phase. The target of a pair was consistently presented at that assigned phase, by which a stable relationship between the association and endogenous theta phase was established. This was achieved by using a closed-loop EEG system that predicted the upcoming mid-frontal theta phase in real-time, allowing the stimulus presentation to be triggered at the desired phase (see Figure 1a).

**Figure 1.**
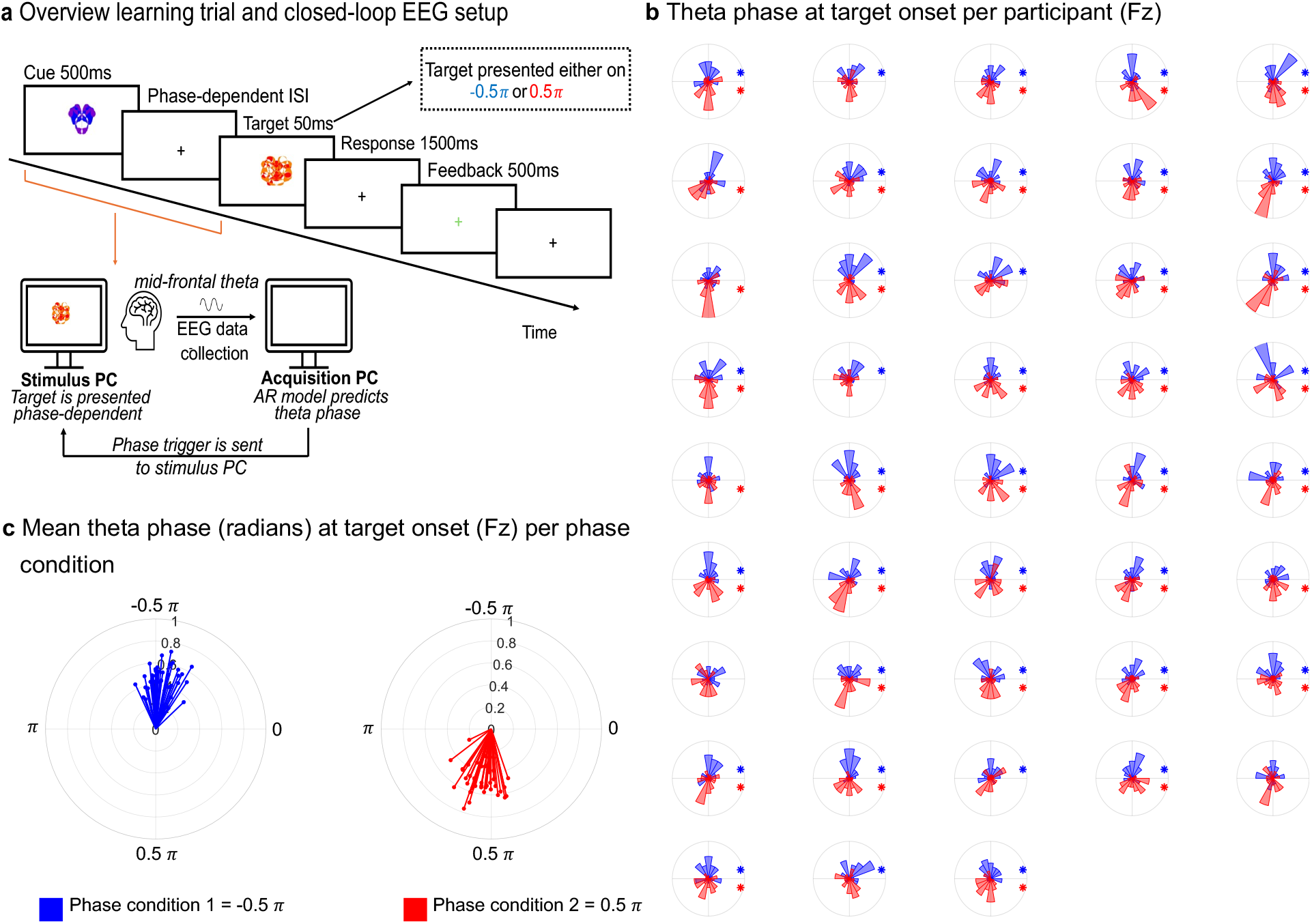
Closed-loop phase-triggered setup. **a)** Schematic overview of one learning trial and closed-loop EEG setup. EEG data is collected and streamed into the acquisition PC. An autoregressive (AR) model predicts theta phase in real-time, after which a trigger is sent to the stimulus PC and the target is presented in a phase-dependent manner. **b)** Participant-wise circular histograms of theta phase at target onset for both phase conditions. Blue and red correspond to phase condition 1 and 2, respectively. Asterisks indicate significant Rayleigh test after FDR correction. **c)** Mean phase vectors per phase condition, one vector representing mean phase angle for one participant, with vector length reflecting phase consistency.

During the learning period, participants were again presented with cue-target pairs, but now a pair could either correspond to the previously presented matching pairs (in 50% of trials in this period) or form a non-matching pair for which one of the other seven target items was randomly selected (in 50% of trials in this period). Matching targets were presented at their pair’s Learned phase, whereas non-matching targets were presented at the opposite phase (for example, a non-matching target following a cue whose pair was assigned to 0.5π was presented at –0.5π). Participants indicated via button press whether the pair was valid or invalid, and feedback was provided after each response (see Figure 1a).

During the final testing period, in which participants underwent memory testing, the trial structure was identical to that of the learning period, except that no feedback was provided and both matching and non-matching targets were presented equally often at both phases. This means, that the phase on which a target was presented during testing (further to referred to as Tested phase) no longer depended on whether it matched the cue; the target could appear at the phase to which its pair had been assigned, or at the opposite phase, with equal probability. Thus, target presentation during retrieval either re-instated or altered the phase-item relationship established during encoding.

### Target stimuli presented at intended phase

We first aimed to validate if our closed-loop approach correctly presented the stimuli at the intended theta phase for each phase condition and participant. To estimate this without contamination from a target-evoked response, we used so called “cue-only” trials in which we presented a cue, but no target, that we implemented in 15.8% of the trials in learning period and in 13.5% of the trials in testing period (see Methods). In these trials, the phase-prediction algorithm still sent a digital trigger at the predicted phase, providing a stimulus-free time-locking point across all tested phases. We first asked if the triggers were reliably phase locked in each condition by running a Rayleigh tests. The majority of participants showed a significant non-uniformity for each phase condition (74 out of 86 FDR corrected significant Rayleigh tests; Figure 1b and 1c), suggesting successful target phase-locking was implemented by our closed-loop EEG system. Note that for the actual presentation of visual targets, there was an additional delay from phase prediction up to target presentation due to the refresh of the screen. We could verify this was kept to a minimum (on average 11.49ms, translating into a phase lag of approximately 0.09π –0.18π across the 4 – 8Hz theta band; see Supplementary Figure S2 for the distributions and Supplementary Methods and Results). Across participants, the average resultant vector length and mean angle for the –0.5π phase condition was *α*ˉ = 1.4587, R = 0.4671 (SD = 0.1114), and *α*ˉ = –1.7779, R = 0.4829 (SD = 0.1179) for the 0.5π phase condition. Additionally, we asked if the triggers in the two phase conditions were locked to different phases. For each participant, we compared phase clustering between the two conditions using a Watson-Williams test. The two conditions differed significantly in all 43 participants (all p < .001; F statistics and degrees of freedom per participant are reported in Supplementary Table S1).

### Increased theta power over Fz before target onset

For phase-specific prediction to be meaningful, the pre-target interval must contain a genuine theta oscillation rather than merely aperiodic activity. We therefore confirmed that the pre-target interval at Fz was characterized by a genuine increase in mid-frontal theta power. We computed the power spectra from the pre-target window of –1 to 0 seconds (s) relative to target onset across all trials in which a target was presented, across all experimental periods. To isolate the periodic component from aperiodic background activity, we parametrized each participant’s power spectrum at Fz using *specparam*, previously *fooof* ^32,33^, and subtracted the estimated aperiodic component from it, which yields a periodic power spectrum. The grand-average periodic spectrum is visualized in Figure 2a, left panel, and shows a clear theta peak around 7Hz. Although a distinct alpha peak was also present, *specparam* resolved theta and alpha as separate oscillations, and we focus on theta here as it formed the basis for our real-time prediction. We additionally visualized the grand-average scalp topography of relative theta power, which we defined here as the ratio of average 4 – 8Hz power to average low-frequency 2–10Hz power at each electrode (Figure 2a, right panel). To visualize the individual participant scalp map (Figure 2b), relative theta topographies were additionally normalized to Fz by dividing the relative theta value at each electrode by the corresponding participant-specific value at Fz. Visual inspection showed that theta activity is not globally elevated, but specifically enhanced in frontal-central regions, relative to neighboring frequencies (Figure 2a and 2b).

**Figure 2.**
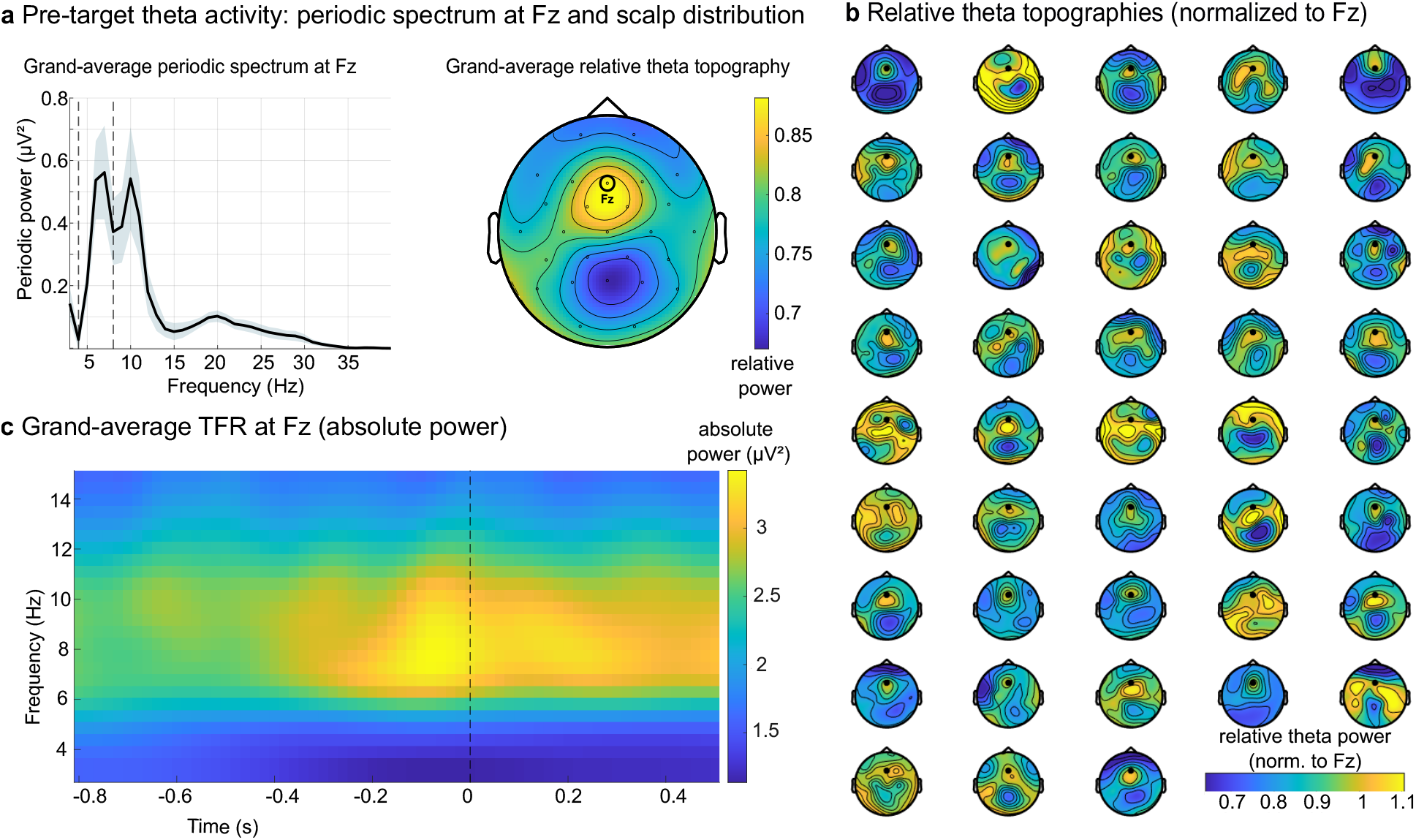
Theta power results. **a)** Grand-average periodic spectrum reconstructed using *specparam* with vertical dotted lines representing theta band of interest, 4 – 8Hz (left panel) and grand-average relative theta topography **b)** Relative theta topographies per participant additionally normalized to Fz. **c)** Grand average time-frequency representation at electrode Fz. Power increases as time point 0 approaches. Note that there is no separate theta and alpha peak as in *a* due to the lower spectral resolution of the time-frequency compared to the frequency analysis in *a*.

Having established that the presence of pre-target theta at Fz, we next asked whether this theta activity was modulated over time around the anticipated target. To investigate the temporal evolution of the theta increase, we ran a time-frequency analysis on the “cue-only” trials. Visual inspection of the time window around the phase-predicted trigger at Fz showed an increase of absolute theta power before the trigger (Figure 2c). To quantify this, we compared theta power (4 – 8Hz) between a baseline window (−0.75 to −0.25 s) and a window straddling the target onset trigger (−0.25 to 0.25s). Theta power was significantly higher in the trigger window compared to the baseline window (mean [trigger window, baseline window] = [2.6331 µV², 2.3383 µV²]; mean difference = 0.295 µV², 95% CI = [0.1710, 0.4190]), t(42) = 4.81, p < .001, which was replicated by a non-parametric Wilcoxon signed-rank test (p < .001). Together, these findings confirm that mid-frontal theta activity increased in anticipation of the target, supporting the assumption underlying our real-time, phase-based prediction approach.

## Behavioral results

### Encoding at 0.5π associated with higher memory sensitivity

Having established the neural validity of our phase-prediction approach, we next turned to the behavioral data to test whether the phase of theta influenced episodic memory formation. Using trials from the testing period of the experiment, we calculated the effect of our experimental manipulations on memory performance. Memory performance was quantified using the sensitivity index (d′), which reflects how well participants distinguished valid, previously learned cue–target pairs from invalid pairs, taking both correct recognitions and false alarms into account. Rather than calculating d′ separately for each condition, we estimated it from the trial-level responses using a Generalized Linear Mixed-Effects Model ^34^ (GLMM) with a probit link, following a signal detection theory framework ^35^. Trials were coded according to whether they contained an invalid pair (Target = 0) or a valid learned pair (Target = 1). With the response dependent variable coding for valid = 1 and invalid = 0, the model estimates the difference in the probability of endorsing valid versus invalid pairs. With a probit link, this difference is expressed on the same standardized scale used to calculate d′, such that the Target coefficient corresponds to d′. At the reference levels of Learned Phase and Tested Phase, the Target coefficient gives the baseline d′, while interactions between Target and the phase predictors indicate whether d′ changes across phase conditions (see Methods). Our final model was the following:

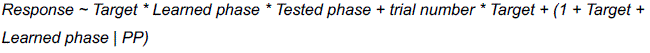

(for the model refinement approach, see Methods). The analysis revealed significant main effects for Trial number (β = –0.0021, standard error (SE) = 0.0004, 95% CI = [-0.0028 –0.0014], p < 0.001), Target (β = 1.5777, SE = 0.1343, 95% CI = 1.3143 1.841], p < 0.001), and Learned phase (β = –0.1468, SE = 0.0723, 95% CI = [-0.2885 –0.0051], p = .0424). The effect of trial number reflects a change in response tendency over time, with participants becoming more conservative (responding “non-match”) over the time course of the experiment, irrespective of accuracy. The significant effect of Target indicates that, at the reference phase conditions, responses differed between valid and invalid trials. This is consistent with above chance discrimination. The significant effect of Learned phase represents the effect of encoding phase on the probability of a “matching” response for the reference target condition (invalid trials) and Tested phase. This indicates a shift in response tendency. Further, we found a significant interaction for Learned phase * Target (β = 0.2629, SE = 0.0933, 95% CI = [0.08 0.4458], p = 0.0049), reflecting the difference in d’ between the two encoding conditions. The positive Learned phase * Target interaction indicates higher d’ for the 0.5π encoding condition relative to the –0.5π reference condition. Thus, memory sensitivity was higher when targets were encoded at 0.5π compared with –0.5π (see Figure 3a). We also found a significant Trial number * Target interaction (β = 0.0024, SE = 0.0005, 95% CI = [0.0014 0.0034], p < 0.001), indicating participants got better over the course of the experiment. We found no significant interaction effects for Tested phase * Target (β = 0.0708, SE = 0.0919, 95% CI = [-0.1093 0.2509], p = 0.4411) and Learned phase * Tested phase * Target (β = –0.1721, SE = 0.1318, 95% CI = [-0.4306 0.0863], p = 0.1917). To show the observed data underlying the non-significant three-way interaction, we visualized the raw memory performance for each combination of Learned phase and Tested phase in Supplementary Figure S3, in which no systematic pattern consistent with the representation account is visible. This is in line with the non-significant interaction within our model. The full model coefficients, model fit, and covariance parameter results are provided in Supplementary Material Table S3.

**Figure 3.**
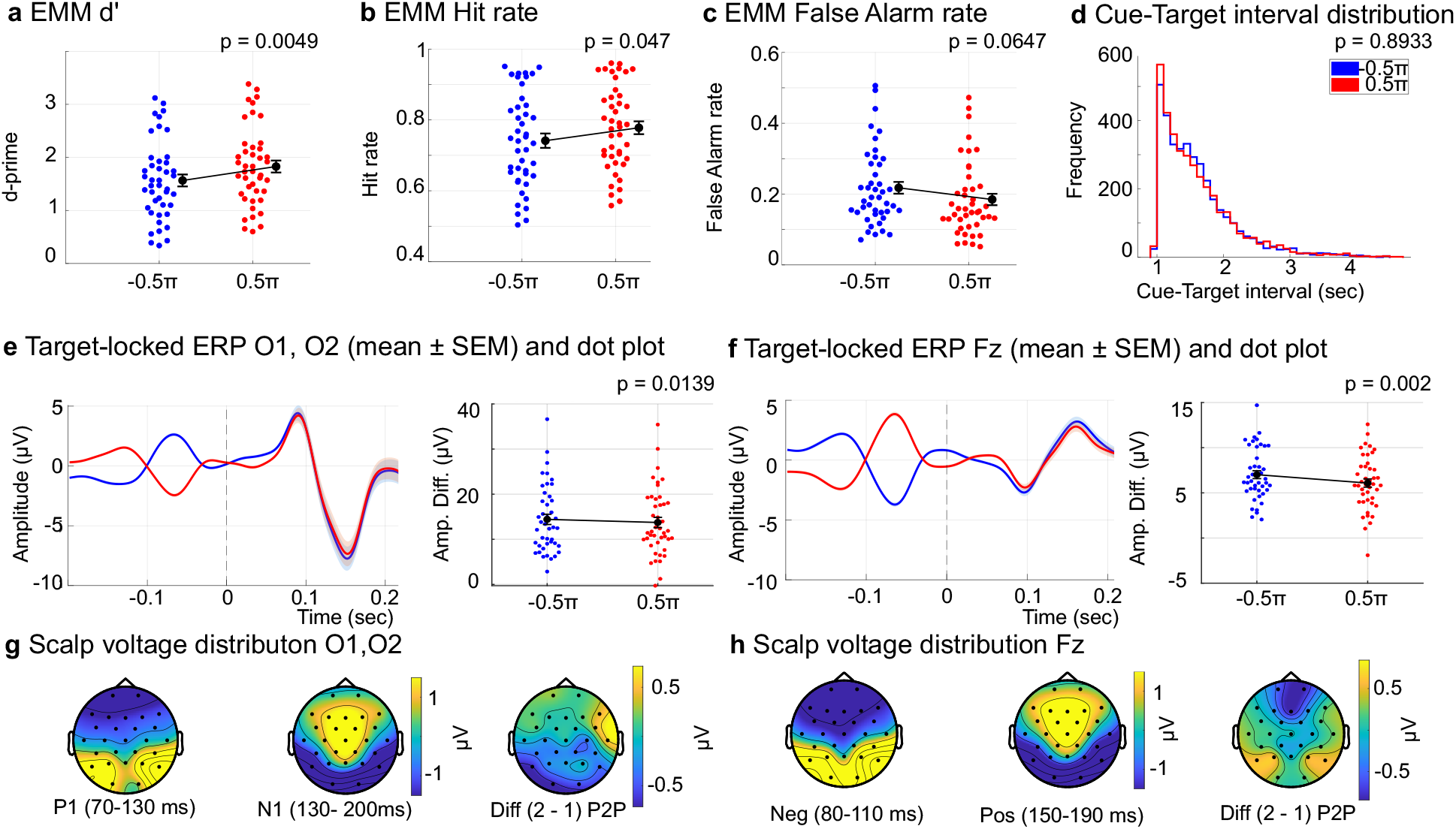
Behavioral results and follow up ERP analysis. Scatter plots depicting the estimated marginal means from the GLMMs, one dot representing one participant. Blue represents Learned phase condition –0.5π; red dots represent Learned phase condition 0.5π. ERPs and scalp voltage distributions over occipital electrodes and Fz. Balack dot = group mean, error bars represent SEM. **a)** Estimated marginal means (EMM) d’ values plotted against Learned phase. Note that the slopes are the same as d’ as the model does not have a random slope. **b)** EMM Hit rates plotted against Learned phase. **c)** EMM False alarm rates plotted against Learned phase. **d)** Distributions of the cue-target interval durations during the learning period of the experiment plotted separately for phase condition –0.5π (blue) and phase condition 0.5π (red). **e** ERP per phase condition over the occipital electrodes (left) and mean peak-to-peak values per participant and phase condition (right). **f)** ERP per phase condition over the frontal electrode Fz (left) and mean peak-to-peak values per participant and phase condition (right). **g)** O1, O2: Collapsed topographies of the ERP and the phase condition difference plot. Left and middle: scalp voltage distributions averaged across both phase conditions within P1 and N1 ERP component windows, respectively. Time windows were selected based on N1 and P1 window. Right: difference in peak-to-peak amplitude between phase conditions across electrodes. **h)** Fz: Collapsed topographies of the ERP components and the phase condition difference plot. Left and middle: scalp voltage distributions averaged across both phase conditions within the early negative and late positive ERP component windows, respectively. Time windows were selected based on the peak latencies observed in the grand-average ERP at Fz. Right: difference in peak-to-peak amplitude between phase conditions across electrodes. Topographies shown in **g** and **h** are for visualization purposes only; the statistical comparisons were performed on the a priori electrodes O1/O2 and Fz.

### Learned phase condition 0.5π associated with increased hit rates and reduces false alarm rate

To follow up on the d’ analysis, we analyzed the effects of our experimental manipulation on hit rates and false alarm rates by fitting separate GLMMs (the full models can be found in Supplementary Tables S2). The GLMM for hit rates showed a significant effect of Learned phase (β = 0.219, SE = 0.1102, 95% CI = [0.0029 0.435], p = 0.047), in line with the d’ results. The positively signed coefficient indicates that participants showed an increase in performance in correctly identifying valid trials when the target was encoded on phase condition 0.5π compared to phase condition –0.5π (see Figure 3b). The remaining effects were not significant (full model coefficients, model fit, and covariance parameter results are provided in Supplementary Material Table 4). For the GLMM of false alarm rates, we found a non-significant trend of Learned phase on false alarm rates (β = –0.2363, SE = 0.1279, 95% CI = [-0.487 0.0144], p = 0.0647), although the estimate suggested lower false alarm rates in the 0.5π condition (see Figure 3c). This finding is also in line with the d’ results. We also find a significant effect of trial number (β = –0.0035, SE = 0.0006, 95% CI = [-0.0047 –0.0022], p < 0.001), indicating that false alarm rate goes down with increasing exposure to the stimuli along the time course of the experiment. The remaining effects were not significant. The full model coefficients, model fit, and covariance parameter results are provided in Supplementary Material Table S5.

### Response times did not differ significantly between phase conditions

To investigate response times, we fitted a separate GLMM (Supplementary Tables S2). Response times significantly decreased over trials, as indicated by a significant main effect of the covariate trial number (β = –0.0005, SE = 0.00004, 95% CI = [-0.0006, –0.0004), p < 0.001, corresponding to a decrease of approximately 0.05% per trial. This suggests that participants responded faster as the task progressed. Our experimental conditions Learned phase and Tested phase did not significantly alter participants response times, indicating that there was no evidence that Learned phase or Tested phase affected response times. The full model coefficients, model fit, and covariance parameter results are provided in Supplementary Material Table S6.

### No evidence that cue-target interval account for the d’ effect

One potential concern is that phase-dependent triggering could have introduced systematic differences in the cue-target interval between the two phase conditions. If the time interval systematically differed between the two phase conditions, any effect on d’ could reflect a difference in timing rather than in endogenous theta phase per se. Because the behavioral effect of phase on d’ emerged from the learning period, we focused this control analysis on those trials. To this end, we fitted a GLMM to predict the effective cue-target time interval dependent on phase condition across all trials. This analysis provided no evidence for a systematic difference in the cue-target interval between phase conditions (β = –0.0017, SE = 0.0127, 95% CI = [-0.0265, 0.0231), p = 0.8933]. Figure 3d shows the distributions of the cue-target interval durations during the learning period of the experiment separately for phase condition –0.5π (blue) and phase condition 0.5π (red). The distributions closely overlap and the estimate difference was near zero, making it unlikely that differences in cue-target interval duration account for the effect of Learned phase on memory sensitivity. The full model coefficients, model fit, and covariance parameter results are provided in Supplementary Material Table S7.

### Sensory processing is unlikely to explain the d’ effect

Next, we considered the possibility that the effect of Learned phase on memory sensitivity could arise from increased sensory processing at that phase, rather than from a memory-related phase preference. To test this, we examined the visual-evoked responses over the occipital electrodes (O1 and O2) and the mid-frontal ERP at Fz, which we have selected a priori. If increased sensory processing could explain the effect of Learned phase on memory sensitivity, we would expect enhanced sensory-evoked responses at those electrodes at the same phase for which learning was improved (phase condition 0.5π).

We first extracted the P1-N1 peak-to-peak amplitude for target stimuli over all learning trials in each electrode. When comparing P1-N1 over the occipital electrodes, we found a significant stronger visual evoked response for phase condition –0.5π (mean = 14.4167 µV) compared to phase condition 0.5π (mean = 13.7318 µV) with t(42) = –2.5686, p = 0.0139, Cohen’s d = –0.3917 and a 95% CI of the phase difference of [-1.2229 –0.1468] (Figure 3e).

Thus, the enhanced sensory-evoked response during encoding was not observed in the condition previously identified as yielding superior memory performance. Scalp voltage distributions averaged across both phase conditions showed a posterior positivity during the P1 time window (70–130ms). In contrast, the N1 (130–200ms) exhibited a broader scalp distribution, extending more anteriorly than the P1. The Phase 2 − Phase 1 peak-to-peak difference map illustrates the electrode-wise distribution of phase-related changes in peak-to-peak amplitude across the scalp and is presented for descriptive purposes (see Figure 3g).

Similarly, when looking at the early negative – subsequent positive component peak-to-peak amplitude over electrode Fz, we found a significantly (t(42) = –3.2937, p = 0.002, Cohen’s d = –0.5023, 95% CI = [-1.4943 –0.3588]) stronger evoked potential for phase condition –0.5π (mean = 7.013 µV) compared to phase condition 0.5π (mean = 6.0864 µV, see Figure 3f).

Descriptively, the early negative component scalp distribution was characterized by a focal posterior positivity, whereas the subsequent positive component scalp distribution was broader and extended toward more anterior scalp regions. The electrode-wise Phase 2 − Phase 1 peak-to-peak difference map showed that changes in P1–N1 amplitude were distributed across posterior and adjacent scalp regions rather than being confined to a single focal area (see Figure 3h). Hence, the visual evoked responses were stronger in the phase conditions with weaker memory performance.

Together, the dissociation between the phase of maximal sensory response and the phase of the behavioral benefit indicates that the behavioral effect at 0.5π is not readily explained by enhanced early sensory proceessing. Instead, the findings are more consistent with a direct influence of theta phase on memory processing.

## Discussion

The current study investigated whether the phase of ongoing mid-frontal oscillations during memory encoding and retrieval influences associative memory performance. We found that memory sensitivity depended on the theta phase on which a target of an associative cue-target pair was encoded: associations for which the target was encoded at 0.5π were remembered better than those encoded at −0.5π. We found no evidence that the phase at retrieval or matching encoding-retrieval phases affected memory sensitivity. Our results thus provide evidence that theta phase modulates encoding efficacy. This is in line with a *gating* account in which phase acts on the process of encoding and by that influences when items are more successfully encoded in memory.

Our finding of phase-dependent encoding fits within a growing number of studies showing that oscillatory phase shapes how incoming information is processed and, in turn, influences behavioural outcomes ^5,6,9,36^. This principle is already better established across perceptual and attentional tasks ^5,6,9,36^, but has also increasingly been demonstrated for mnemonic processes ^13,37^. Our findings add to this picture by showing that the ongoing mid-frontal theta phase at encoding shapes how well an association is later remembered. Encoding efficacy therefore does not appear to be constant, but rather fluctuates with the ongoing mid-frontal theta rhythm. Importantly, by targeting endogenous theta phase in real time, our study goes beyond post-hoc associations between EEG phase and memory, providing more direct evidence that the ongoing theta state at encoding contributes to subsequent memory success.

We found that prefrontal theta phase during encoding influences associative binding. Successful associative binding depends on a network primarily involving prefrontal cortex and hippocampus, with the hippocampus supporting the formation of associations between elements of an experience and prefrontal regions contributing to the organization and control of encoding, particularly when mnemonic demands are high ^38,39^. Theta oscillations were proposed as a mechanism organizing the communication between regions within this network ^39,40^, with mid-frontal theta reflecting the frontal node activity ^10,31,41^. Presenting stimuli at different phases of this mid-frontal theta might have placed them in a network state that is more or less favorable for associative binding, leading to improved or reduced memory performance, respectively.

Because the EEG scalp measured phase does not directly correspond to a physiological state ^42^, the effect we observe in the current study reflects a difference in encoding efficacy between the two phases separated by π. Recent evidence illustrated that single-neuron activity in the human hippocampus is coordinated with frontal theta during working memory, with higher phase locking being observed when memory load is high ^43^. Although this research is not testing associative memory, it shows how frontal theta structures the timing of hippocampal processing^43^. In this sense, theta phase can be seen as an internal process through which the memory system organizes encoding.

Considering that previous perceptual and attention studies also report phase dependent effects, one might wonder whether our findings could be a consequence of phase-dependent sensory processing or attentional sampling rather than from the process of memory encoding itself ^44,45^. Indeed, enhanced sensory processing at specific theta phases could improve the quality of the information available for encoding. Similarly, rhythmic attentional sampling could produce phase-dependent fluctuations in perceptual gain, resulting in better memory for stimuli presented at phases of heightened attention. Both enhanced sensory and attentional processes have typically been associated with increased ERP amplitudes, especially in the early sensory components P1 and N1 ^46,47^. Hence, if differences in sensory processing or attention would explain the behavioral effect, we would have expected increased ERP responses in the phase condition that also showed the effect for increased memory performance. However, we found a dissociation between the phase of superior behavioral performance and largest target-evoked sensory responses: The theta phase optimal for memory performance did not coincide with the phase condition that showed enhanced early sensory responses. Why the sensory-evoked response was enhanced at one particular phase cannot be determined from our current experimental design and is not critical for our interpretation. We do not take this dissociation to show that sensory processing was less effective at the memory-favorable phase, but rather as arguing against a simple account in which stronger sensory or attentional responses drive better memory. This is for two reasons: First, stronger neuronal responses do not always equate greater information content available ^48,49^. Second, the relationship between ERP amplitude at encoding and subsequent memory itself is not straightforward as successful memory has been associated with different ERP amplitude patterns depending on the component and encoding conditions ^50,51^. Finally, if our results were merely explained by increased sensory processing or attention, one would have also expected an effect of retrieval phase on memory performance. We found no evidence for such effect. Taken together, the combination of behavioral evidence, ERP evidence, and the absence of evidence for an effect of retrieval phase on memory performance support an encoding specific role of theta phase in associative memory, though the precise neuronal mechanism remains open.

We found no support for the phase as *representation* account, which would have predicted that memory performance is increased whenever the phase of a target at testing matches the phase on which the target was encoded. We found no evidence that the externally measured theta phase at encoding functioned as a retrievable feature of the memory representation. The lack of support for phase as a retrievable component of the memory representation does not mean that theta oscillations are not involved in organizing information at different theta phases as proposed by the theta-gamma code hypothesis. According to this hypothesis, different neuronal groups coding for different neuronal representations have preferred firing phases that are organized along the theta cycle ^20^. Instead, our current findings suggest that theta phase is likely not a necessary component of the memory representation; rather, phase may determine the temporal organization of neuronal firing during encoding, while the resulting pattern of neuronal activity constitutes the memory representation. In other words, the absence of increased memory performance whenever encoding and retrieval phase match in the current study does not contradict the idea that theta phase contributes to the formation of memory representations. In our current study, we find no evidence that externally measured phase at which an item is encoded functions as a feature that can be externally imposed and subsequently reinstated to facilitate retrieval.

Another influential account for the role of theta phase in memory is the Separate Phase for Encoding and Retrieval (SPEAR) model ^15,52^. This model proposes that memory encoding and retrieval are optimized at different phases of the hippocampal theta cycle, thereby minimizing interference between these two competing processes ^15,52^. Indeed, it has been observed that episodic memory encoding fluctuates in a theta rhythm ^13^. Additionally, it was shown that the optimal phase for memory retrieval is separated by roughly half a cycle relative to the optimal phase for memory encoding along the theta oscillation ^53^. In the present study, we found no evidence for a segregation between optimal encoding and retrieval phase: memory performance only varied with encoding, and not with retrieval phase. Importantly, the SPEAR model specifically concerns hippocampal theta ^15,52^, whereas in the current study we measured mid-frontal theta. Even though the prefrontal cortex and the hippocampus form a functional network, their rhythms are not necessarily always coupled ^54,55^, and additionally, hippocampal theta has been reported at lower frequencies than the theta band targeted here ^11^. Our findings hence do not provide a direct test of the phase-specificity of encoding and retrieval as proposed by the SPEAR model. Nevertheless, the encoding effect may still be in line with the model.

SPEAR proposes that during encoding, external input to the hippocampus is strong and weak during retrieval, where internally driven recall dominates instead. Because our manipulation acted on externally presented stimuli, whose influence the model locates at encoding, an effect of encoding rather than retrieval phase is consistent with the model ^15^. Future studies should thus test the SPEAR model directly by directly targeting hippocampal theta oscillations as a closed-loop target, for example by using MEG ^56^.

The closed-loop EEG setup of the current study enabled prospective control over endogenous theta oscillations. By using this set-up, we gained experimental control over the phase at stimulus presentation during encoding and retrieval, allowing us to directly test the influence of phase during encoding and retrieval on memory performance. Very few studies have tested the memory effects of endogenous theta phase. Examples come from intracranial recordings ^57^ and MEG/EEG studies ^19,53^, but did not actively manipulate the phase at which items were encoded or retrieved. While previous closed-loop approaches have already demonstrated that neural activity can be monitored and used prospectively to time stimulus presentation in different domains ^58–60^ and brain stimulation applications ^61,62^, the current study is the first one to apply a closed-loop EEG approach to probe the phase of endogenous mid-frontal theta during memory encoding and retrieval. This extends previous applications of closed-loop approaches to the phase of endogenous theta oscillations, which in turn provides the functional means to test the relevance of theta phase for memory.

The methodological advantage of this approach is relying on reliably tracking an oscillatory band and using it to prospectively target certain phases of the ongoing rhythm for stimulus presentation. We have validated that this was implemented successfully with our current approach: we showed that the oscillatory signal we leveraged contained a periodic theta-band component, we find evidence for an increase in theta activity in the 1s window before target onset, and most importantly, we showed reliable phase-locking of target stimuli, showing that the system presented the targets at the desired phase. Future approaches could improve the current closed-loop approach further, for example by reducing the closed-loop latency, applying a stricter power threshold (which does, however come at the cost of a lower amount of trials), or by isolating the periodic component of the oscillatory signal before the online phase estimation ^32^.

Using a closed-loop EEG approach, we here show that the mid-frontal theta phase on which targets of associative pairs were encoded affected how well these were subsequently remembered. This pattern is consistent with a *gating* account, in which theta phase acts on the process of encoding. These findings indicate that the rhythmic state during the moment of encoding shapes what information is later remembered better. Building on these findings, future work incorporating methods sensitive to hippocampal activity could investigate how this phase-dependent modulation of encoding emerges from interactions within the broader frontal– hippocampal memory network.

## Methods

### Participants

We tested N = 48 participants (N = 34 female, mean age (SD) in years = 21,42 (± 2.63)). Participants were recruited from the Faculty of Psychology and Neuroscience (FPN) at Maastricht University (UM) via flyers and an online recruitment system ^63^. Participants were instructed to arrive at the laboratory well-rested and free from the influence of any intoxicants. Written informed consent was given prior to participation, and participation was reimbursed either monetarily (7.50 Euros per hour) or by course credits. The current study was approved by the Ethics Review Committee Psychology and Neuroscience (approval number OZL_264_26_02_2023) and was additionally reviewed by the local EEG project proposal committee at UM, FPN. The project followed the safety procedures proposed by UM, FPN.

### Closed-Loop EEG

EEG data were recorded with a sampling rate of 10000Hz at a sub-sample divisor of 20 and a nominal sampling rate of 500Hz, using the actiCHamp EEG ^64^. We positioned 28 electrodes on the scalp of the participant using an international 10-20 system EEG cap ^65^: O1, O2, P7, P3, Pz, P4, P8, CP5, CP1, CP2, CP6, T7, C3, Cz, C4, T8, FC5, FC1, FC2, FC6, F7, F3, Fz, F4, F8, Fp1, Fp2, A2, A1. AFz was the ground electrode. Three additional electrodes, lateral to both eyes and vertically below the left eye, were used to monitor eye movements. The reference electrode for online referencing was A2. All electrodes were positioned using the Nuprep Skin Prep Gel and Signa Gel Electrode Gel. Electrode impedance was kept below 10kΩ during the whole session, which was assessed using the BrainVision recorder software ^66^. The EEG signal was streamed to the lab-streaming layer (LSL) with the actiCHamp Connector ^67^.

### Real-time EEG processing

The aim of the current study was to systematically present the target of an associative cue-target pair at one out of two possible theta-phases, measured at Fz. To this end, we presented a visual target dependent on ongoing mid-frontal theta phase (see Figure 1). EEG data were streamed into the data acquisition PC, and a signal processing pipeline was carried out using custom code in MATLAB 2023b ^68^. Using an autoregressive (AR) model, the EEG signal was predicted in real-time in the time domain ^61,69^. This prediction is necessary because of the inherent technical delay in the system as well as edge artifacts introduced by bandpass filtering used for estimating phase (but also see ^70^ for a bandpass free method for phase prediction). We implemented our approach by following the procedures used by Shirinpour et al. (2020) but changed parameters for our study design as we saw fit. Specifically, the electrode of interest was Fz, and we applied a band-pass filter to the signal between 4 and 8Hz, focusing on the target frequency *tf* = 6 ± 2Hz. The window length for online processing amounted to *window length = sampling frequency*3/mean(tf)* (sampling frequency further to reffered as *fs*). The desired target phase, two different phases at which a target should be presented, were defined as –0.5*π and 0.5*π. We chose these phases to have comparable oscillatory amplitudes. We additionally accounted for a technical delay of 10ms. This accounted for delays in recording and sending of the EEG signal, the streaming of the signal from the LSL to the data acquisition PC and online processing in MATLAB. The AR model was initialized with several parameters: *AR order* was set to as *fs*2/mean(tf)*. *AR edge,* defined as the number of samples to be removed from the beginning and end of each data segment (accounting for the edge artifact), was set to *fs*0.2/mean(tf),* corresponding to 34ms. *Forward samples,* defined as the number of future samples the AR model predicts, was set to *fs*1/mean(tf)*. Lastly, a *power threshold* was applied to ensure that phase information was only used when a theta oscillation of sufficient amplitude was present, as phase-prediction accuracy declines when oscillatory power is low ^61^. The relevant power was computed as the squared amplitude of the analytic signal (Hilbert transform) of the band-pass-filtered (4–8 Hz), AR-predicted signal at the electrode of interest, taken at the predicted target sample. The threshold was set to an absolute value of 1 µV², applied identically across participants, and was determined heuristically based on inspection of pilot data, chosen so that phase prediction remained reliable while retaining a sufficient number of trials. A target was only triggered when the power exceeded this threshold.

To present the target based on the desired phase, each incoming EEG chunk was pulled from the LSL buffer and re-referenced to the common average of all active EEG electrodes. For the phase analysis, the signal at the electrode of interest (Fz) was then extracted and band-pass filtered to the theta band (4–8 Hz) using a finite impulse response (FIR) filter. To ensure that the signal is suitable for further processing, and not contaminated by eye-movement artifacts, we conducted an analysis of the vertical electrode channel (VEOG). To this end, data from the VEOG channel was pulled from the LSL buffer for the specified window length and demeaned. If any data point in VEOG exceeded 100 µV, the signal was considered to contain an eye blink and was not used for phase prediction. In these cases, no target trigger was sent, and the phase prediction was resumed once there was no artifact detected anymore. To predict the EEG signal, AR model coefficients were estimated using the Yule-Walker method ^71,72^, implemented using MATLAB’S *aryule* function ^73^. Once the signal was predicted, the Hilbert transform was applied to extract the instantaneous phase and power of the signal. As soon as the phase was forward-predicted to be close to the desired phase and the power threshold of 1 µV^2^ was met, a trigger was sent to the stimulus PC, which then presented the target based on the desired phase. The maximum possible target delay was set to 5 s (with a 0.5 s buffer). On a small number of trials, the target phase was not reached within this window: 0 of 689 passive trials (0.00%), 23 of 8,969 learning trials (0.26%), and 20 of 9,546 testing trials (0.21%). These trials were excluded from analysis.

Next to a digital trigger that logged cue and target stimulus onset, we also recorded the precice stimulus onset with a photodiode. The photodiode was placed in the upper left corner of the stimulus PC to not interfere with the participants vision, and a small rectangular patch of the display underneath the photodiode turned black concurrently with the stimulus being presented on the screen.

### Associative memory task

The task design of the current associative memory task, including stimuli, was taken from van de Ven et al., 2017 ^30^. Participants completed an associative memory task, for which they learned eight pairs of abstract visual stimuli. For every participant, stimuli were randomly drawn out of a list containing eight possible cues and eight possible targets. Cues and targets were abstract, non-conceptual stimuli generated using Fourier descriptors, with cue stimuli colored in blue and purple and target stimuli colored in red and orange (size = 2.57° x 3.42°, Supplementary Figure S1). The task consisted of three periods: a passive exposure period, a learning period, and a testing period. Throughout the experiment, participants were instructed to avoid head movements and minimize eye blinks as much as possible to ensure good EEG data quality.

### Passive exposure period

During the passive exposure period, participants were shown all eight stimulus pairs twice and were encouraged to memorize them. Each trial started with the presentation of the cue, displayed for 500ms, followed by a black fixation cross for an inter-stimulus interval (ISI) contingent on phase prediction (see section Real-time EEG processing), with a minimal ISI of 0.5s and a maximal ISI of 4.5s. The target was then presented for 50ms, after which the black fixation cross reappeared. The inter-trial interval (ITI) was randomly jittered between 3 and 4.5s, indicating the interval between target onset of the previous trial and cue onset of the next trial. Critically, four of the eight targets were randomly assiged to be presented at the desired phase of –0.5π, and the other four targets were presented at the desired phase of 0.5π. To prime for and encourage rapid responses in the following phases, participants were instructed to press the key “m” immediately after the target presentation. The maximum allowed response time was 1.5s, and responses were logged for offline analysis. If a response was given in time, the fixation cross turned grey. If participants did not respond within 1.5s, a message appeared on the screen for 500ms, informing them their response was too slow before moving on to the next trial.

### Learning period

During the learning period, participants strengthened their memory of the stimulus pairs. The general trial structure as well as parameters were kept the same as in the passive exposure period. In each learning block, each cue was represented four times in a randomized order. After the fixation cross, a probe was presented for 50ms, after which the black fixation cross reappeared (Figure 1A). In 50% of trials, the probe was the target initially associated with the cue, while in the other 50% of trials, the probe was a non-target drawn from the seven non-target alternatives. Participants were required to judge the validity of each stimulus pair, pressing “m” for a valid pair (where the probe was the target, “match”) and “n” for an invalid pair (where the probe was a non-target, “non-match”). Importantly, when the probe was a target, it was always presented at the same desired theta phase as in the passive exposure period, while non-targets were presented at the opposite pahse condition. Feedback was provided after each response: a green fixation cross (displayed for 500ms) indicated a correct judgment, and a red fixation cross (displayed for 500ms) indicated an incorrect judgment. The maximum allowed response time was 1.5s, with responses logged for offline analysis. If participants did not respond within 1.5s, a message appeared on the screen for 500ms, informing them their response was too slow, before moving on to the next trial. On average, participants faild to repond within the response time on mean = 6.91 trials (SD = 5.62) across all task periods and trials. Participants completed a up to eight blocks, each containing 32 cue-target trials, or until they reached a performance accuracy of at least 75%. Participants completed a minimum of 2 and a maximum of 8 learning blocks (mean = 5.48, SD = 1.94). Each block also included six trials in which a cue purposefully was not followed by a target. These “cue-only” trials were designed to estimate the accuracy of phase prediction without the confounding influence of event-related potentials (ERPs) elicited by the target event. In these trials, the fixation cross turned blue to signal that no response was required.

### Testing period

During the testing period, participants completed several test trials following a similar procedure to that of the learning period, with the same trial structure and parameters as in the learning period, apart from two crucial differences. First, participants did not receive any feedback on their performance; the fixation cross turned grey after a response was given. Second, the association between the matching target of a pair and the phase that was established throughout the passive exposure and learning period was broken: The target was either presented at the learned phase, or at the phase with which the target had not been associated with, with a probability of 0.5. Presenting the target of a matching pair either on it’s associated phase or on its non-associated phase is the crucial manipulation for our study design in order to test the temporal context account proposed for the role of theta phase in memory. Under this assumption, we expect that memory performance is enhanced when an item is retrieved at the same phase as it was encoded ^9,19,20^. Therefore, our study design is a 2 x 2 factorial design, with independent variables being the Learned phase (–0.5π, 0.5π) and the Tested phase (–0.5π, 0.5π). In total, participants completed three blocks with 74 trials each, comprising 16 trials in each task condition, plus ten “cue-only” trials where no response was required.

## Analysis

### Participant exclusion criteria

A chi-square test was carried out in MATLAB 2023b ^68^ to assess if individual performance does not differ significantly from chance performance. For this analysis, all trials from the testing period were pooled and analyzed. Participants were excluded if there was either no evidence for performance significantly differently from chance level, or if performance accuracy amounted to 100% in all conditions. We excluded N = 4 based on no significant deviation from chance level performance. We excluded N = 1 based on 100% performance accuracy in all conditions. Our sample size therefore amounted to N = 43 participants. On average, participants completed 5.49 learning blocks (SD = 1.94).

## EEG analysis

### Epoching

#### Stimulus onset determination

The stimulus onset sample was defined using the photodiode deflection that we recorded from the display throughout the experiment. In “cue-only” trials, no photodiode signal was recorded as no physical diode event was happening in these cases.

Nevertheless, the appearance of the fixation cross was marked by a digital trigger sent to the EEG system. We therefore relied on the digital system trigger timestamps for epoching of “cue-only” trials.

#### Epoch time window

For the purpose of pre-processing, continuous EEG data of all experimental periods were segmented into epochs time-locked to cue-onset. Epochs started 1s before cue-onset and extended until 2s after the target onset. Trials without a valid target onset trigger were excluded. Additionally excluded was the final trial of each experimental block since we observed systematic recording artifacts at block termination. All remaining valid epochs were retained for subsequent preprocessing analysis, and epochs were either re-epoched or had a new 0 point defined dependent on the subsequent analysis.

### Preprocessing

EEG data of all experimental periods were preprocessed and analyzed in MATLAB ^68^ using fieldtrip ^74^. After Discrete-Fourier-Transform based line noise removal, data were detrended and demeaned. Scalp electrode channels were visually and spectrally inspected, and noisy channels were manually rejected. Data were then re-referenced to the average of remaining EEG channels. Independent component analysis (ICA) was performed and primarily used to identify eye-blinks and horizontal eye movements ^75^. We identified components based on topography, time course, and spectral characteristics and removed them subsequently. The few rejected scalp EEG channels (on average 0.05, SD = 0.3, ranging from 0-2 channels across participants) were reconstructed using weighted interpolation based on neighboring electrodes. Neighbors were identified using triangulation of the electrode layout, and interpolation weights were computed from the spatial proximity of surrounding channels. Finally, residual artifacts were removed by visual inspection of single trials using fieldtrip’s *ft_rejectvisual* in summary mode. Candidate outliers were identified interactively based on the summary statistics provided by the software and removed without applying a pre-defined threshold. All pre-processing decisions were stored in participant specific-log files. On average, we rejected 19.74 trials (SD = 20.88) from cue-locked epochs, reflecting a mean percentage rejected trials of 5.52% (SD = 4,72%). The maximum percentage of rejected trials in these epochs is 22.17%. After pre-processing, the number of retained trials ranged in between 256 and 530 for all participants (mean = 416.38, SD = 70.69). We did not exclude trials based on behavioral performance, as our primary objective of the EEG analysis was to assess the effectiveness of the methodological setup across all trials.

### Phase analysis: Validating intended phase matches instantaneous phase during target presentation

This analysis included “cue-only” trials from all experimental periods; trials in which we purposefully did not present a target after a cue: After the cue, AR-based phase estimation still took place, and a system trigger was sent to the acquisition PC, but no target was presented to the participant. This allowed us to estimate target phase without target-elicited ERP confounds in the signal. Offline analysis included only the signal from Fz (which was used for phase estimation). The initially cue-locked epochs were recentered, such that target onset corresponded to time point zero. We used the same electrode and temporal alignment approach for the frequency analysis as well as the time-frequency analysis. We included a minimum of 20 and a maximum of 46 trials (mean = 29.79, 95% CI = [27.98 31.60]) for phase condition –0.5π and a minimum of 14 and a maximum of 45 trials (mean = 29.37, 95% CI = [27.22 31.53]) for phase condition 0.5π per participant for the phase analysis.

To ensure consistency with our real-time phase extraction during the experiment, the same procedure for phase estimation was applied in the validation analysis: The signal was first bandpass filtered in the theta range (4 – 8Hz) using a two-pass FIR filter. We then extracted instantaneous phase using the Hilbert transform. For each trial, we extracted instantaneous phase at target onset. Values were represented as unit vectors on the complex plane, which were then averaged within each participant and condition to obtain a mean phase vector. To assess if target onset occurred at a preferred theta phase for each phase condition, we used Rayleigh test for non-uniformity using the CircStat toolbox in MATLAB ^76^. For each condition and participant, which resulted in 86 separate Rayleigh tests, we computed the resultant vector length, Rayleigh test statistic, and p-value. We used False Discovery Rate (FDR) correction to correct for multiple comparisons. Subsequently, we assessed for each participant if the two phase conditions were significantly different from each other using Watson-Williams tests in CircStat toolbox in MATLAB ^76^. The test was applied to the previously calculated mean phase angles and resultant vector lengths.

### Frequency analysis: Validating theta power over Fz

EEG data were analyzed at our electrode of interest, Fz. We included all trials in which a target was presented from all experimental periods. We did not differentiate between phase condition –0.5π and 0.5π because an increase in power is needed throughout the whole experiment to ensure precise phase estimation, irrespective of phase condition. Per participant, we included a minimum of 222 and a maximum of 452 trials (mean = 357.19, 95% CI = [338.92 375.45]). We selected a pre-target time window of –1 to 0s relative to target onset for the spectral analysis.

Power spectra were computed using a Fast-Fourier Transform approach (*mtmfft*) with a Hanning taper. To verify that the pre-target theta increase reflected a genuine oscillation rather than a shift in broadband (aperiodic) activity, we parameterized the Fz power spectra using the *specparam* algorithm ^32^ (formerly *fooof*; called from MATLAB via the *fooof_mat* wrapper). For this analysis, power spectra were estimated over a broader 2–40 Hz range to allow a robust estimate of the aperiodic component. The model was fit separately for each participant on their trial-averaged Fz spectrum, using a knee aperiodic mode and a peak width limit of 2–12 Hz. For one participant whose spectrum did not converge under the knee model, the aperiodic component was fit using the fixed mode instead. For each participant, the aperiodic component estimated by the model was subtracted from the power spectrum, yielding a periodic (aperiodic-corrected) spectrum expressed in absolute power (µV²/Hz). These periodic spectra were then averaged across participants, and the SE of the mean across participants was computed at each frequency to visualize the group-level oscillatory spectrum.

To characterize the scalp distribution of pre-target theta activity, we additionally computed subject-level theta topographies. For each electrode, relative theta power was defined as the average power across the 4–8 Hz range divided by the mean power across the 2–10 Hz range, with power estimated per trial and averaged across trials within each participant, hence expressing theta activity at each electrode relative to the participant’s overall low-frequency power in the pre-target interval. Participant-level topographies were grand-averaged to obtain the group-level scalp distribution of relative theta activity. To additionally visualize the consistency of this distribution across participants, individual topographies were normalized to Fz by dividing each electrode’s relative theta value by the corresponding value at Fz, such that values express relative theta power at each electrode proportional to that at Fz.

### Time-Frequency analysis: Validating theta increase over Fz right before target onset

Trials for the time-frequency and phase analysis were restricted to “cue-only” trials, so that these subsequent analyses would not be influenced by target-elicited ERPs, and both phase-conditions were pooled together. Across participants, we included a minimum of 34 and a maximum of 78 trials (mean = 59.16, 95% CI = [55.63 62.70]). Time-frequency representations of oscillatory power were computed using a sliding time-window approach (*mtmconvol*) with a Hanning taper. Power was estimated from 3 – 20Hz in steps of 0.5, using a frequency-dependent window length of 4 cycles per frequency. Estimates were obtained for time points from – 1s before target onset and 1safter target onset in steps of 20ms. To reduce edge artifacts, data were zero-padded to the next power of two, and mirror padding was used. Power estimates were averaged across trials per participant.

To test if theta power significantly increased before target onset, we compared theta power (4 – 8Hz) at Fz between a baseline window (–0.75 and –0.25s relative to target onset) and a window straddling target onset (–0.25 and 0.25s). The two windows were defined to be non-overlapping, and the shared boundary was assigned to the baseline window. For each participant, theta power was averaged across the 4 –8Hz range and across all time points within each window, resulting in one value per window and per participant. These values were then compared in a paired-samples t-test. To confirm that the result did not depend on the assumptions of the parametric test, we additionally ran a non-parametric Wilcoxon signed-rank test. Because the frequency dependent analysis window at the lower theta frequencies (1000ms at 4Hz) exceeds the 500ms comparison windows, power estimated near the window boundary integrated activity from both sides; this temporal smearing renders the comparison conservative, such that a significant difference between the two windows indicates the effect is present despite this smoothing.

## Behavioral analysis

### Trial wise exclusion criteria

Unless stated otherwise, the behavioral analysis only included trials from the testing period and trials were excluded when no response from the participant was given. This included cases when participants pressed the wrong button, if the maximum prediction time or maximum response time was reached, or “cue-only” trials. After removing these trials, we identified response time outliers separately for each participant using the interquartile range (IQR) method which provides a robust approach for the skewed distribution of reaction time data, while at the same time accounting for individual differences in response speed. We removed trials that fell below Q1 – 1.5 * IQR or above Q3 + 1.5 * IQR, where IQR = Q3 – Q1. The number of included trials per participant ranged from 176 to 190 (95% CI = [181.66, 184.15]. Out of an initial total number of trials of 9546, we excluded 1452 (15,2%) based on no response logged (comprising cue-only trials, in which no response was required by design, trials with no valid participant response, or trials in which maximum prediction time was met). We excluded an additional 229 (2,4%) based on the IQR filtering. We remained with a total number of trials of 7865.

## Statistical analysis

Analyses were conducted in MATLAB 2023b. We calculated the d’ sensitivity index, hit rates, false alarm rates, and response times for each participant. Statistical analyses were conducted using MATLAB’s *fitGLME* function to account for the repeated measures structure of the data.

All GLMMs were estimated using maximum likelihood with the Laplace approximation. Fixed effects were based on our a-priori hypothesis, which are described below. We first included a random intercept to account for individual differences in overall memory performance in a base model. We then evaluated random slopes for the within-participant predictors, prioritizing predictors that showed the strongest fixed effects in the base model, as these effects were of particular interest and could plausibly vary across participants. Random slopes were added sequentially, with each model compared with the preceding model using MATLAB’s *compare* function. The final random-effects structure was determined based on model fit and convergence. We did not include interactions of the random slopes in any model to avoid unreliable beta estimates. The threshold for statistical significance was set at p < .05.

### D’ sensitivity index

To analyze the d’ sensitivity index for each participant, we followed the approach described by Heynckes et al. (2020). Following ^35^, d’ was estimated directly with the GLMM framework. The full model was specified as:

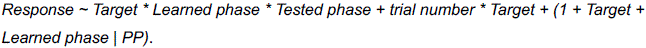

The behavioral outcome (Response) reflects whether participants indicated that a valid pair was presented on a given trial. This binary outcome is modeled as a function of the predictor variable Target, coding for the actual presence or absence of a valid pair. Valid (“match”) trials are treated as signal-present (hits), and invalid (“non-match”) trials are treated as signal-absent (false alarms). A probit link function and binomial distribution are used to fit the model, aligning the model with an equal-variance Gaussian signal detection theory framework. Binomial size was set to one, reflecting single-trial observations. Categorical predictors were reference coded, with −0.5π as the reference level for Learned phase and −0.5π as the reference level for Tested phase. With the probit link, the coefficient for Target represents d′ at the reference levels of Learned phase and Tested phase. The interactions with Target allow d′ to vary across the different Learned phase and Tested phase conditions. This modelling framework allows the inclusion of fixed and random effects to test whether d’ varies systematically across Learned phase, Tested phase, and their interaction. Trial number (centered to its mean) was included as a covariate to account for potential changes in performance over the course of the experiment, such as practice or learning effects.

We included Target, Learned phase, Tested phase and their interaction as fixed effects in the model since we were interested in the effect of Learned phase and Tested phase on memory sensitivity, d’. The interaction between trial number and Target was included to allow sensitivity (i.e., the difference between signal and noise responses) to change over the course of the experiment. Random effects were added incrementally using an additive, nested model comparison approach. All models included a by-subject random intercept. Random slopes were then introduced stepwise, starting with predictors that showed the strongest fixed effects (Target, Learned phase), followed by Tested phase. When the tested slope significantly improved the model fit, the slope was retained. The final model can be found in Table S1.

#### Hit rate, False alarm rate

To analyze hit rates and false alarm rates as a follow up analysis of the d’ results, we fitted separate GLMM with a binomial distribution and logit link function. For the hit-rate analysis, only signal-present (match) trials were included, whereas for the false-alarm-rate analysis, only signal-absent (non-match) trials were included. Each model included the fixed effects Learned phase, Tested phase, their interaction. Trial number (centered) was included as a covariate to account for potential changes in performance over the course of the experiment. A random intercept was included for each participant to model subject-level baseline variability. Random slopes were added incrementally, and model fits were compared. Random slopes for Learned phase and Tested phase were evaluated sequentially using nested model comparisons. A random slope was retained when its inclusion significantly improved model fit. The final model per outcome measure can be found in Table S1.

#### Response times

Response times (in seconds) were calculated relative to target onset for trials with a recorded response, irrespective of response accuracy. To analyze response times, we fitted a GLMM using a gamma distribution and a log link function. The model included the fixed effects Learned phase, Tested phase, their interaction, as well as trial number (centered). The model included a random intercept for each participant. Random slopes for Tested phase and Learned phase were subsequently evaluated using nested model comparisons. The final model per outcome measure can be found in Table S1.

### Follow up analysis: GLMM of cue-target interval duration imposed by closed-loop EEG setup

Because the phase-dependent presentation of the target was controlled by the closed-loop EEG system, we performed a post-hoc control analysis to verify that this procedure did not introduce systematic difference in the cue-target interval durations between the phase conditions. This might otherwise confound behavioral effects. Because we found a significant effect of Learned phase on d’ in the current study, we focused our analysis on trials belonging to the Learning period of the experiment. For each trial, cue-target interval duration was calculated as the time between cue and target photodiode onset. Trials with a recorded response time of zero were excluded because these trials corresponded to “cue-only” trials or trials without a recorded response. No IQR filtering was applied here as the purpose of this analysis was to characterize the timing of the closed-loop stimulus-presentation procedure rather than behavioral performance. We analyzed the interval durations in MATLAB 2023b by using GLMM with MATLAB’s *fitGLME* function, accounting for repeated measures across participants. The model included Phase (phase at which the target was presented during the learning period) and centered trial number as fixed effects. We included a random intercept for each participant; and subsequently introduced Phase as a random slope. The slope was retained since it significantly improved model fit. The final model can be found in Table S1. We used a gamma distribution and log link function to account for the positively right-skewed cue-target interval distribution.

### Follow up analysis: Event-related potentials (ERPs) analysis over occipital and frontal electrodes after target onset

As an exploratory follow-up analysis, we examined whether Learned phase was associated with differences in target-evoked neural responses that could contribute to the observed phase-dependent behavioral effect. Given the significant effect of Learned phase on d′, we focused this analysis on trials from the Learning period of the experiment. First, we performed an analysis on visual evoked responses. Target-locked epochs were low-pass filtered at 30Hz using a two-pass FIR filter with a Hamming window. Baseline correction was applied using the –0.2 to 0s interval relative to target onset. Trials were sorted based on phase condition. For each participant and phase condition, ERPs were computed for all channels by averaging across trials with fieldtrip’s *ft_timelockanalysis*. Then, to quantify differences between visual evoked responses, we focused our analysis on the occipital channels, which we have selected a priori because we expect visual evoked response at this site to be maximal. ERPs were averaged across electrode O1 and O2 for each participant and phase condition. The two time-windows of interest were defined by using the P1 and N1 component, with an interval from 70-130ms and 130 to 200ms, respectively. For each participant, peak-to-peak amplitude was computed as the difference between the maximum amplitude in the P1 window and the minimum amplitude in the N1 window. We focused on peak-to-peak amplitude differences as we have per definition different baselines in our phase conditions. Differences between the conditions were assessed using a paired samples t-test comparing peak-to-peak amplitudes between the phase conditions. Significance was evaluated at the group level across participants.

In addition to our visual evoked response analysis, we performed a target-locked ERP analysis at our electrode of interest, Fz, which we also selected a priori. To use fixed time windows across participants and conditions, the ERP time windows were determined from the grand-average waveform collapsed across experimental conditions. The average waveform was inspected to determine the typical latencies of the early negative and subsequent positive peaks, yielding analysis windows of 80–110ms and 150–190ms, respectively. These fixed windows were then applied in the subsequent peak-to-peak analyses of individual participant data. For each participant and phase condition, we calculated peak-to-peak differences and performed significance testing in the same way as for the occipital channels. The number of trials per participant ranged from 24 to 124 in phase condition 1 (mean = 81.95, 95% CI = [72.98 90.93]) and from 24 to 124 in phase condition 2 (mean = 81.70, 95% CI = [72.63 90.77]).

