## Supplementary Materials for "Theta-Phase Targeted Encoding Gates Memory Sensitivity"

### Supplementary Methods

#### *Validation of digital system trigger use in "cue-only" trials*

We wanted to investigate the delay between the digital trigger and the actual stimulus presentation trigger, as determined by a photodiode positioned on the upper left corner of the stimulus PC. A small patch located below the photodiode changed the contrast at stimulus onset, which allows for the precise detection of the physical stimulus onset time. We calculated the milliseconds (ms) passed in between digital trigger and respective photodiode onset. Analysis of the descriptive statistics include trials from all three experimental periods of all participants. "Cue-only" trials were not included.

### Supplementary Results

#### *Physical stimulus timing matched the expected display refresh rate*

System trigger-to-photodiode delays were highly similar for cue and target stimuli. For cue events, the mean delay was 11.4922ms (SD = 6.3707), with a minimum of 4ms and a maximum of 40ms. The median delay was 10ms, and the interquartile range (IQR) was 10ms (25h percentile 6ms, 75$^{th}$ percentile = 16ms). Importantly, 95% of cases showed delays below 22ms. For target events, the mean trigger-to-diode delay was 11.5187ms (SD = 6.4982ms), also with a minimum of 4ms and a maximum of 40ms. The median was again 10ms, with an IQR of 10ms (25$^{th}$ percentile = 6ms, 75$^{th}$ percentile = 16ms). For visualization of the distributions, see Supplementary Figure S2. Hence, overall, both cue and target timing distributions were narrow and comparable, which indicates a consistent correspondence between digital system triggers and the actual physical stimulus onset as measured by the photodiode. Because the monitor operated at 60Hz (one refresh = 16.67ms), delays in the range of one refresh cycle were expected. Consistent with frame-locked presentation, most trials fell within a single refresh interval, with no evidence of systematic multi-frame delays.

### Supplementary Figures

**Figure S1**

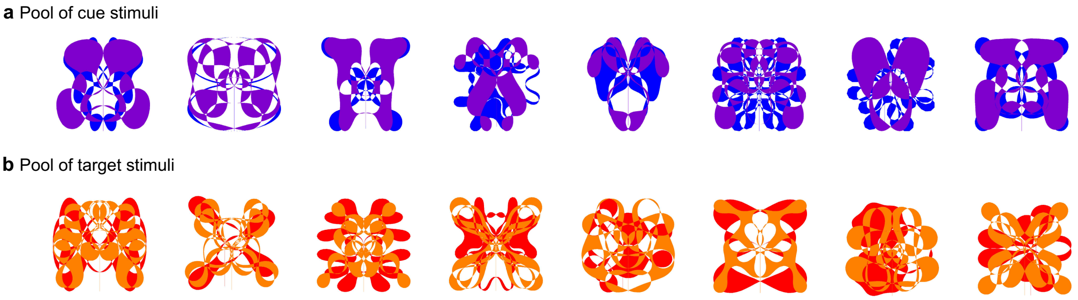


Abstract visual cues (purple-colored abstract patterns) and visual targets (orange -colored abstract patterns) used in the experiment. Eight pairs consisting of a cue and target were drawn for each participant randomly.

**Figure S2 Onset delays**

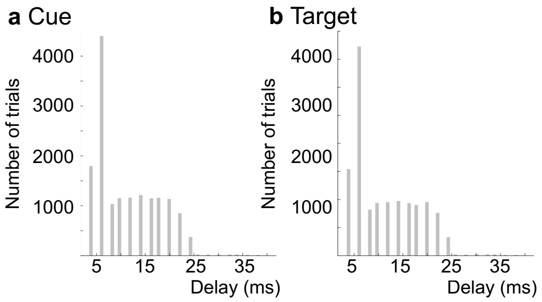


Distribution of stimulus onset delays in between the photodiode deflection and digital trigger plotted for cue (a) and target (b) appearances separately. Here, we show all trials from all experimental phases of all participants. Trials considered are the same as for behavioral analysis.

**Figure S3 Observed d' by Learned phase and Tested phase**

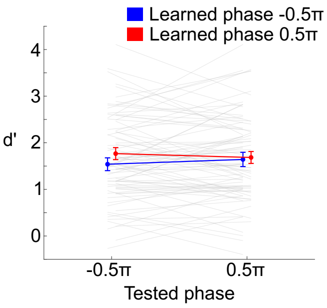

Raw (observed) d’ for each combination of Learned phase and Tested phase. In contrast to the main text figures, which report EMMs from the GLMM, we here display raw (observed) directly. We computed d’ directly from each participant’s observed hit and false alarm rates for every combination of Learned phase and Tested phase. Group means (±SEM) are displayed in color, whereas grey lines mark individual participants. The plot shows no systematic pattern that is consistent with the reinstatement hypothesis – increased d’ whenever the Learned phase matches the Tested phase. This is consistent with the non-significant interaction between Target*Learned phase*Tested phase reported in the main text.

### Supplementary Tables

**Table S1. Per-participant Watson-William test results**

| PP | -0.5π | 0.5π | WW F statistic | df_between | df_within | WW_p-value |
|---|---|---|---|---|---|---|
| PP6 | 25 | 26 | 51.1495 | 1 | 49 | < 0.001 |
| PP7 | 30 | 31 | 34.9294 | 1 | 59 | < 0.001 |
| PP8 | 39 | 39 | 72.3567 | 1 | 76 | < 0.001 |
| PP9 | 27 | 19 | 42.3054 | 1 | 44 | < 0.001 |
| PP11 | 23 | 29 | 45.0012 | 1 | 50 | < 0.001 |
| PP12 | 29 | 28 | 29.1753 | 1 | 55 | < 0.001 |
| PP13 | 26 | 24 | 53.5598 | 1 | 48 | < 0.001 |
| PP14 | 39 | 39 | 58.3551 | 1 | 76 | < 0.001 |
| PP15 | 28 | 26 | 44.8422 | 1 | 52 | < 0.001 |
| PP17 | 27 | 19 | 98.9404 | 1 | 44 | < 0.001 |
| PP18 | 30 | 22 | 37.212 | 1 | 50 | < 0.001 |
| PP19 | 24 | 22 | 70.3649 | 1 | 44 | < 0.001 |
| PP20 | 26 | 21 | 33.3514 | 1 | 45 | < 0.001 |
| PP21 | 41 | 29 | 53.822 | 1 | 68 | < 0.001 |
| PP22 | 23 | 24 | 45.0016 | 1 | 45 | < 0.001 |
| PP23 | 26 | 26 | 79.853 | 1 | 50 | < 0.001 |
| PP24 | 25 | 38 | 22.2551 | 1 | 61 | < 0.001 |
| PP25 | 34 | 34 | 75.4776 | 1 | 66 | < 0.001 |
| PP26 | 33 | 32 | 72.6435 | 1 | 63 | < 0.001 |
| PP27 | 22 | 29 | 44.14 | 1 | 49 | < 0.001 |
| PP28 | 32 | 32 | 34.9695 | 1 | 62 | < 0.001 |
| PP29 | 21 | 25 | 78.0594 | 1 | 44 | < 0.001 |

| | | | | | | |
|---|---|---|---|---|---|---|
| PP30 | 29 | 30 | 70.1656 | 1 | 57 | < 0.001 |
| PP31 | 27 | 23 | 37.2542 | 1 | 48 | < 0.001 |
| PP32 | 35 | 34 | 42.3344 | 1 | 67 | < 0.001 |
| PP33 | 36 | 38 | 97.1034 | 1 | 72 | < 0.001 |
| PP35 | 29 | 45 | 58.1245 | 1 | 72 | < 0.001 |
| PP36 | 26 | 23 | 40.9894 | 1 | 47 | < 0.001 |
| PP37 | 32 | 33 | 66.5558 | 1 | 63 | < 0.001 |
| PP39 | 46 | 29 | 34.4713 | 1 | 73 | < 0.001 |
| PP40 | 20 | 14 | 28.363 | 1 | 32 | < 0.001 |
| PP41 | 30 | 29 | 78.937 | 1 | 57 | < 0.001 |
| PP42 | 26 | 25 | 51.2246 | 1 | 49 | < 0.001 |
| PP43 | 36 | 37 | 52.6443 | 1 | 71 | < 0.001 |
| PP44 | 34 | 38 | 53.7392 | 1 | 70 | < 0.001 |
| PP45 | 32 | 33 | 62.6523 | 1 | 63 | < 0.001 |
| PP46 | 38 | 36 | 121.4478 | 1 | 72 | < 0.001 |
| PP47 | 38 | 37 | 42.9303 | 1 | 73 | < 0.001 |
| PP48 | 29 | 28 | 75.0392 | 1 | 55 | < 0.001 |
| PP49 | 21 | 18 | 16.028 | 1 | 37 | < 0.001 |
| PP51 | 28 | 42 | 63.1405 | 1 | 68 | < 0.001 |
| PP52 | 27 | 25 | 38.7506 | 1 | 50 | < 0.001 |
| PP53 | 32 | 32 | 75.7402 | 1 | 62 | < 0.001 |

Note. -0.5π = Phase condition -0.5π, 0.5π = Phase condition 0.5π, WW = Watson-Williams, df = degrees of freedom.

**Table S2. Wilkinson notation of final model in each analysis**

| | |
|---|---|
| D’ sensitivity index | Response ~ Target * Learned phase * Tested phase + trial number * Target + (1 + Target + Learned phase \| PP) |
| Hit rate | Hit ~ Learned phase * Tested phase + trial number + (1 \| PP) |
| False Alarm rate | False Alarm ~ Learned phase * Tested phase + trial number + (1 + Learned phase \| PP) |
| Response time | Response time ~ Learned phase * Tested phase + trial number + (1 \| PP) |

| | |
|---|---|
| Cue-Target interval duration | Cue-Target interval duration ~ Phase * trial number + (1 + Phase \| PP). |

**Table S3. GLMM Results d'**

Fixed effects coefficients

| Name | Estimate | SE | pValue | Lower | Upper |
|---|---|---|---|---|---|
| (Intercept) | -0.8347 | 0.0743 | < 0.001 | -0.9804 | -0.689 |
| Tested phase | -0.055 | 0.0656 | 0.4024 | -0.1837 | 0.0737 |
| Learned phase | -0.1468 | 0.0723 | 0.0424 | -0.2885 | -0.0051 |
| trial number | -0.0021 | 0.0004 | < 0.001 | -0.0028 | -0.0014 |
| Target | 1.5777 | 0.1343 | < 0.001 | 1.3143 | 1.841 |
| Tested phase * Learned phase | 0.1263 | 0.0945 | 0.1815 | -0.059 | 0.3117 |
| Tested phase * Target | 0.0708 | 0.0919 | 0.4411 | -0.1093 | 0.2509 |
| Learned phase * Target | 0.2629 | 0.0933 | 0.0049 | 0.08 | 0.4458 |
| trial number * Target | 0.0024 | 0.0005 | < 0.001 | 0.0014 | 0.0034 |
| Tested phase *Learned phase * Target | -0.1721 | 0.1318 | 0.1917 | -0.4306 | 0.0863 |

Model fit statistics

| AIC | BIC | Log Likelihood | Deviance |
|---|---|---|---|
| 7812 | 7923.5 | -3890 | 7780 |

Random effects covariance parameters

| | | | |
|---|---|---|---|
| Intercept | Intercept | SD | 0.3804 |
| Learned phase | Intercept | Corr | 0.1616 |
| Target | Intercept | Corr | 0.3290 |
| Learned phase | Learned phase | SD | -0.7872 |
| Target | Learned phase | Corr | -0.2688 |

| | | | |
|---|---|---|---|
| Target | Target | SD | 0.7699 |

Note. SE = Standard error. Lower, Upper: Lower and upper boundary of confidence interval. SD = Standard deviation. Corr = correlation.

**Table S4. GLMM results Hit rate**

Fixed effects coefficients

| Name | Estimate | SE | pValue | Lower | Upper |
|---|---|---|---|---|---|
| (Intercept) | 1.2199 | 0.156 | < 0.001 | 0.914 | 1.5258 |
| Tested phase | 0.0245 | 0.1081 | 0.8205 | -0.1874 | 0.2365 |
| Learned phase | 0.219 | 0.1102 | 0.047 | 0.0029 | 0.435 |
| trial number | 0.0005 | 0.0006 | 0.4146 | -0.0007 | 0.0017 |
| Tested phase * Learned phase | -0.0834 | 0.156 | 0.5929 | -0.3893 | 0.2224 |

Model fit statistics

| AIC | BIC | Log Likelihood | Deviance |
|---|---|---|---|
| 18571 | 18609 | -9279.7 | 18559 |

Random effects covariance parameters

| | | | |
|---|---|---|---|
| Intercept | Intercept | SD | 0.88497 |

Note. SE = Standard error. Lower, Upper: Lower and upper boundary of confidence interval. SD = Standard deviation.

**Table S5. GLMM results False Alarm rate**

Fixed effects coefficients

| Name | Estimate | SE | pValue | Lower | Upper |
|---|---|---|---|---|---|
| (Intercept) | -1.3797 | 0.1317 | < 0.001 | -1.6378 | -1.1215 |
| Tested phase | -0.0975 | 0.1142 | 0.3935 | -0.3214 | 0.1265 |

| | | | | | |
|---|---|---|---|---|---|
| Learned phase | -0.2363 | 0.1279 | 0.0647 | -0.487 | 0.0144 |
| trial number | -0.0035 | 0.0006 | < 0.001 | -0.0047 | -0.0022 |
| Tested phase * Learned phase | 0.2199 | 0.1648 | 0.1823 | -0.1032 | 0.543 |

Model fit statistics

| AIC | BIC | Log Likelihood | Deviance |
|---|---|---|---|
| 18764 | 18814 | -9373.8 | 18748 |

Random effects covariance parameters

| | | | |
|---|---|---|---|
| Intercept | Intercept | SD | 0.6792 |
| Learned phase | Intercept | Corr | -0.0807 |
| Learned phase | Learned phase | SD | 0.3290 |

Note. SE = Standard error. Lower, Upper: Lower and upper boundary of confidence interval. SD = Standard deviation. Corr = correlation.

**Table S6. GLMM results response times**

Fixed effects coefficients

| Name | Estimate | SE | pValue | Lower | Upper |
|---|---|---|---|---|---|
| (Intercept) | -0.3375 | 0.0248 | < 0.001 | -0.3862 | -0.2889 |
| Tested phase | 0.0071 | 0.0078 | 0.365 | -0.0082 | 0.0224 |
| Learned phase | 0.004 | 0.0078 | 0.6087 | -0.0113 | 0.0193 |
| trial number | -0.0005 | 0 | < 0.001 | -0.0006 | -0.0004 |
| Tested phase * Learned phase | -0.0086 | 0.0111 | 0.4374 | -0.0303 | 0.0131 |

Model fit statistics

| AIC | BIC | Log Likelihood | Deviance |
|---|---|---|---|
| 423.74 | 472.53 | -204.87 | 409.74 |

Random effects covariance parameters

| | | | |
|---|---|---|---|
| Intercept | Intercept | SD | 0.1588 |

Note. SE = Standard error. Lower, Upper: Lower and upper boundary of confidence interval. SD = Standard deviation.

**Table S7. GLMM results Cue – Target interval duration**

Fixed effects coefficients

| Name | Estimate | SE | pValue | Lower | Upper |
|---|---|---|---|---|---|
| (Intercept) | 0.461 | 0.0225 | 0 | 0.43047 | 0.4915 |
| Phase | -0.0017 | 0.0127 | 0.8933 | -0.0265 | 0.0231 |
| trial number | 0.0001 | 0.0001 | 0.1157 | -0.0002 | 0 |

Model fit statistics

| AIC | BIC | Log Likelihood | Deviance |
|---|---|---|---|
| 4575.2 | 4623.5 | -2280.6 | 4561.2 |

Random effects covariance parameters

| | | | |
|---|---|---|---|
| Intercept | Intercept | SD | 0.0951 |
| Phase | Intercept | Corr | -0.1464 |
| Phase | Phase | SD | 0.0649 |

Note. SE = Standard error. Lower, Upper: Lower and upper boundary of confidence interval. STD = Standard deviation. Corr = correlation.
